# Etiology of fever in Ugandan children: identification of microbial pathogens using metagenomic next-generation sequencing and IDseq, a platform for unbiased metagenomic analysis

**DOI:** 10.1101/385005

**Authors:** Akshaya Ramesh, Sara Nakielny, Jennifer Hsu, Mary Kyohere, Oswald Byaruhanga, Charles de Bourcy, Rebecca Egger, Boris Dimitrov, Yun-Fang Juan, Jonathan Sheu, James Wang, Katrina Kalantar, Charles Langelier, Theodore Ruel, Arthur Mpimbaza, Michael R. Wilson, Philip J. Rosenthal, Joseph L. DeRisi

**Author notes:** These authors contributed equally to this work (Akshaya Ramesh, Sara Nakielny). e-mail address for authors: Akshaya Ramesh; Sara Nakielny; Jennifer Hsu; Mary Kyohere; Oswald Byaruhanga; Charles de Bourcy; Rebecca Egger; Boris Dimitrov; Yun-Fang Juan; Jonathan Sheu; James Wang; Katrina Kalantar; Charles Langelier; Theodore Ruel; Arthur Mpimbaza; Michael R. Wilson; Philip J. Rosenthal; Joseph L. DeRisi.

## Abstract

**Background:** Febrile illness is a major burden in African children, and non-malarial causes of fever are uncertain. We built and employed IDseq, a cloud-based, open access, bioinformatics platform and service to identify microbes from metagenomic next-generation sequencing of tissue samples. In this pilot study, we evaluated blood, nasopharyngeal, and stool specimens from 94 children (aged 2-54 months) with febrile illness admitted to Tororo District Hospital, Uganda.

**Results:** The most common pathogens identified were *Plasmodium falciparum* (51.1% of samples) and parvovirus B19 (4.4%) from blood; human rhinoviruses A and C (40%), respiratory syncytial virus (10%), and human herpesvirus 5 (10%) from nasopharyngeal swabs; and rotavirus A (50% of those with diarrhea) from stool. Among other potential pathogens, we identified one novel orthobunyavirus, tentatively named Nyangole virus, from the blood of a child diagnosed with malaria and pneumonia, and Bwamba orthobunyavirus in the nasopharynx of a child with rash and sepsis. We also identified two novel human rhinovirus C species.

**Conclusions:** This exploratory pilot study demonstrates the utility of mNGS and the IDseq platform for defining the molecular landscape of febrile infectious diseases in resource limited areas. These methods, supported by a robust data analysis and sharing platform, offer a new tool for the surveillance, diagnosis, and ultimately treatment and prevention of infectious diseases.

## Introduction

The evaluation of children with fever is challenging, particularly in the developing world. A febrile child in sub-Saharan Africa may have a mild self-resolving viral infection or may be suffering bacterial sepsis or malaria—major causes of disability and death [1], [2]. In the past, febrile illness in children under five years of age in most of Africa was treated empirically as malaria due to the limited availability of diagnostics and the risk of untreated malaria progressing to life-threatening illness. This strategy changed with new guidelines from the World Health Organization (WHO) in 2010, which recommend limiting malaria therapy to those with a confirmed diagnosis [3]. However, standard recommendations for management of febrile children who do not have malaria are lacking. Increased knowledge about the prevalence of non-malarial pathogens associated with fever is needed to inform management strategies for febrile children who do not have malaria [2].

Advances in genome sequencing hold promise for addressing global infectious disease challenges by enabling unbiased detection of microbial pathogens without requirement for the extensive infrastructure of modern microbiology laboratories [4], [5]. Although sequence-based diagnostics have not yet replaced most traditional microbiological assays, this situation is rapidly changing, as sequence-based strategies are incorporated for clinical care [6]–[10], and costs have declined dramatically over the past decade [11]. A significant roadblock toward implementation of sequence-based diagnostics is the extensive computational requirements and bioinformatics expertise required. In fact, as sequencing costs decline, computational expenses may proportionally increase due to inevitable expansion of existing genomic databases.

To address these challenges, we developed IDseq, a cloud-based open-source bioinformatics platform and service for detection of microbial pathogens from metagenomic next-generation sequencing (mNGS) data. IDseq requires minimal computational hardware and is designed to enhance accessibility and build informatics capacity in resource limited regions. Here, we leveraged IDseq and mNGS data to perform an exploratory proof-of-concept molecular survey of febrile children in rural Uganda to characterize pathogens associated with fever, including both well-recognized and novel causes of illness.

## Analyses/Results

### Clinical presentations of children providing samples for analysis

From October to December, 2013, 94 children admitted to Tororo District Hospital were enrolled (Table 1). Their mean age was 16.4 (IQR: 8.0-12.0) months, and 66 (70.2%) were female. Chief symptoms reported in addition to fever were cough (88.3%), vomiting (56.4%), diarrhea (47.9%), and convulsions (27.7%). Top admitting diagnoses were respiratory tract infection (57.4%), gastroenteritis/diarrhea (29.8%), and septicemia (11.7%) (Data file 1a).

**Table 1:**
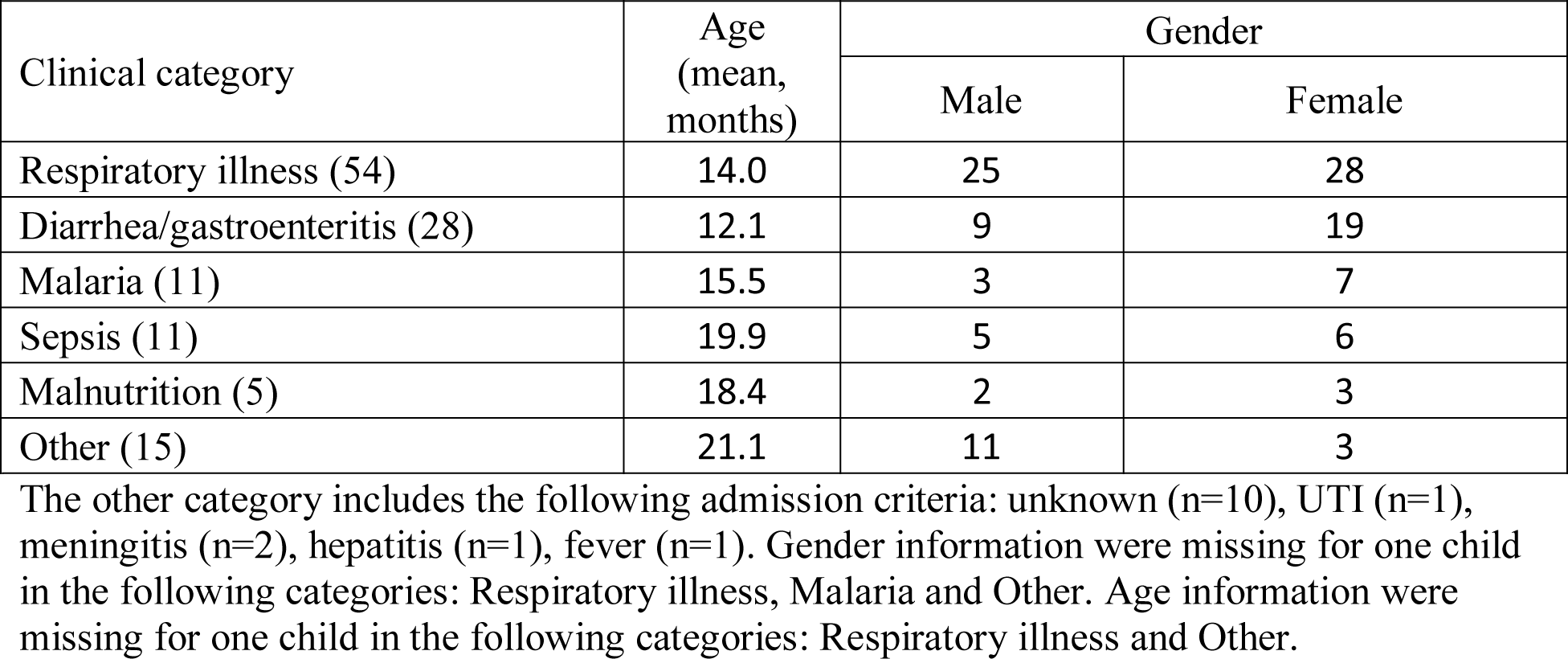
Overview of the patients enrolled in the study

Serum and nasopharyngeal (NP) swab samples were collected from 90 children each; for four children, only one of the two sample types was successfully collected. Although 45 (47.9%) of the children had a presenting symptom of diarrhea, stool samples were available for only 10 due to logistical constraints. Blood smears identified *P. falciparum* in 12 of the 90 samples that underwent mNGS analysis.

### Metagenomic sequencing findings

mNGS was performed on 90 serum, 90 NP swab, and 10 stool samples and analyses was done using the IDseq pipeline (see Methods section, Figure 1); detailed findings are reported in Data file S1b. For each sample type, RNA was extracted, libraries were prepared, samples were sequenced, and reads were analyzed using the IDseq Portal. A mean of 11.5 million (IQR 6.4 – 15.2 million) paired-end reads were obtained per sample; sequencing statistics are in Data file Table S2. For one batch of serum samples, only a single read, rather than paired-end reads, was produced.

**Figure 1:**
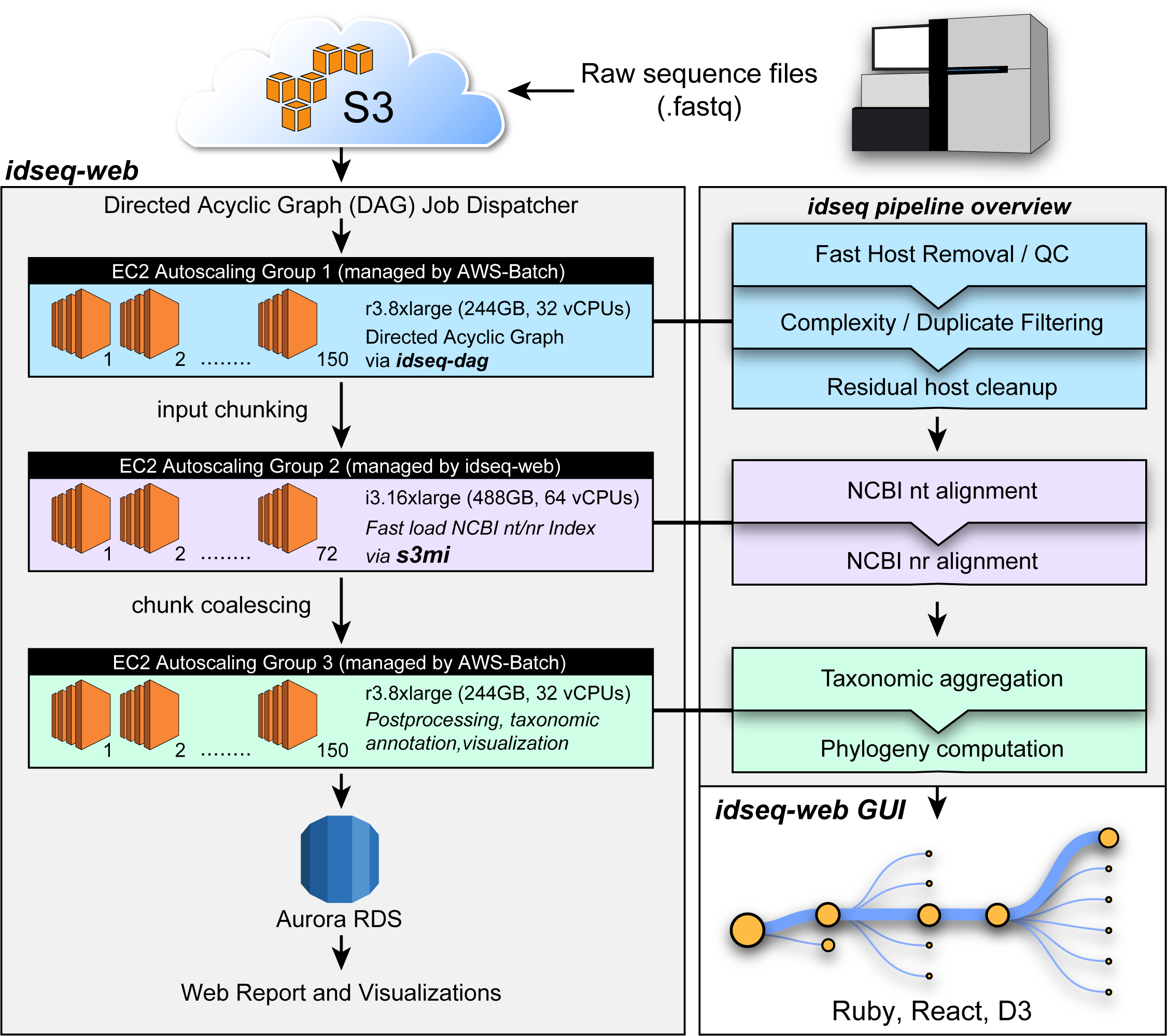
Schematic representation of IDseq pipeline

### mNGS of serum

At least one microbial species was detected in 60 (66.7%) serum samples; more than one microbe was detected in 11 (12.2%) samples (Figure 2A). The most commonly identified microbes were *Plasmodium falciparum* (46, 51.1%) and parvovirus B19 (4, 4.4%). *P. falciparum* was detected in 10/12 samples from patients reported as smear-positive for malaria parasites. mNGS detected *Plasmodium spp.* in 37 additional samples that were smear negative (36 *P. falciparum*, 1 *P. malariae*). Viruses detected from serum included human immunodeficiency 1 virus (HIV-1), hepatitis A virus, rotavirus A, human herpesvirus (HHV) type 6, HHV type 4, HHV type 7, human rhinovirus (HRV)-C, HRV-A, enteroviruses (enterovirus A71, Coxsackievirus B2 and echovirus E30), human parechovirus 2, hepatitis B virus, a novel orthobunyavirus (described in greater detail below), human cardiovirus (Saffold virus), mamastrovirus 1 and Norwalk virus (Figure 2A).

**Figure 2:**
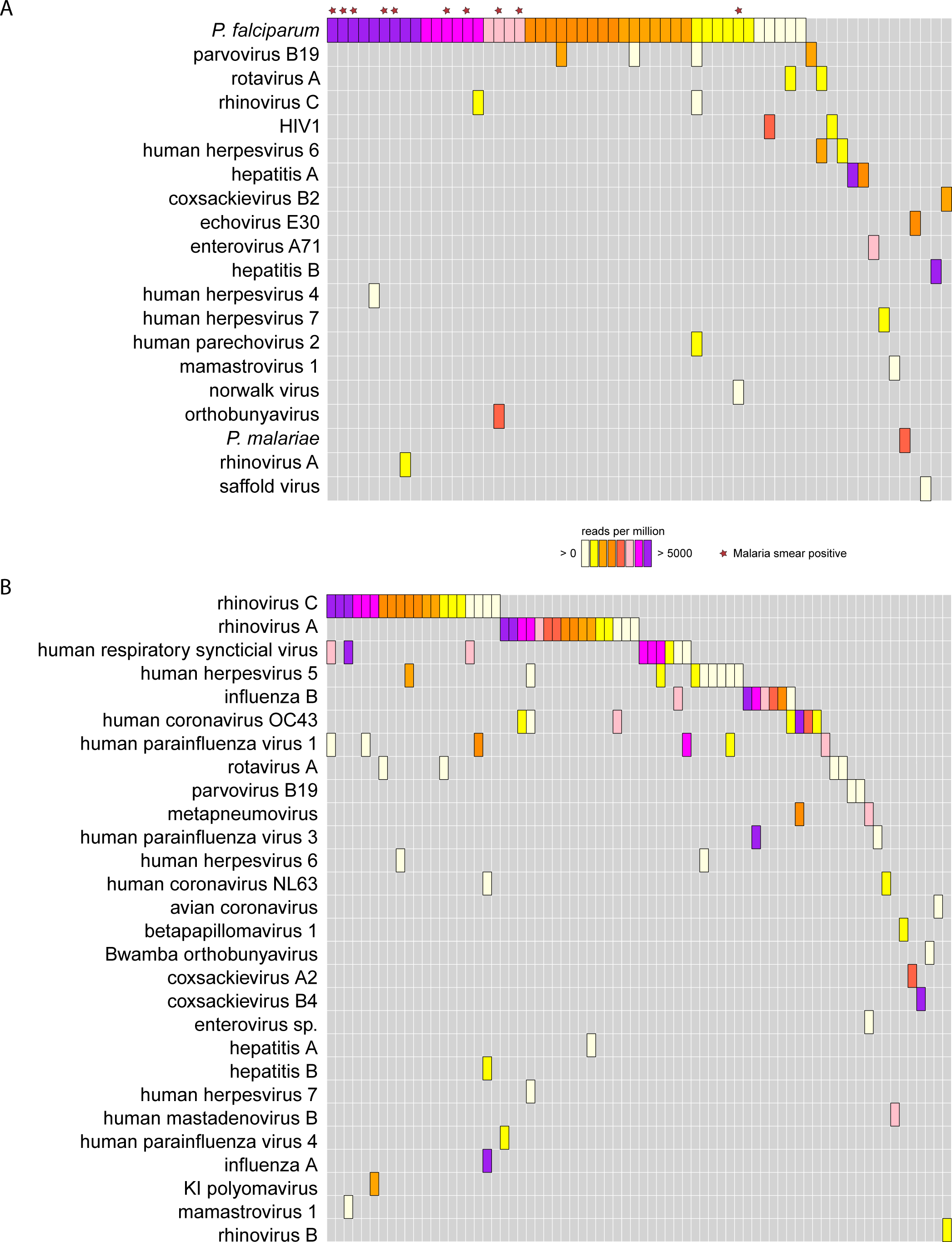
Microbes found in (A) serum and (B) nasopharyngeal (NP) swab samples. Note that bacterial species were not considered for NP samples.

Multiple viruses were detected from serum in patients with *Plasmodium* infections (10 of 46 (21.7%) samples; Table S1a). Three of the four identified parvovirus B19 infections were associated with *P. falciparum*. Additionally, GB virus C and torque teno virus (TTV), which are of unknown clinical significance [12], [13], were identified in the serum of 25 (27.8%) and 37 (41.1%) children, respectively.

### mNGS of NP swabs

A total of 90 NP swabs was collected and processed; 52 (57.7%) of these were from patients with admission diagnoses of pneumonia, respiratory tract infection, or bronchiolitis (Table 1). Chest imaging was not available to further assess these diagnoses. 73 NP samples (81.1%) contained at least 1 viral species (Figure 2B). HRV-A and HRV-C were the most prevalent, followed by respiratory syncytial virus (RSV), cytomegalovirus (HHV-5), influenza B, and coronavirus OC43. Other respiratory viruses identified included influenza A, HRV-B, adenovirus B, 3 human parainfluenza virus types, metapneumovirus, coronavirus NL63, avian coronavirus, coxsackievirus A2, coxsackievirus B2, polyomaviruses (KI), HHV-6 and HHV-7. Other viruses identified that are not typically considered respiratory pathogens included hepatitis A virus, hepatitis B virus, parvovirus B19, mamastrovirus 1, Bwamba orthobunyavirus, betapapillomavirus 1, and rotavirus. Additionally, TTV was found in 49 (54.4%) NP swab samples, including one sample with both gemykrogvirus and TTV. For 26 (28.8%) patients, mNGS identified respiratory viral co-infections, most commonly with HRV-C (n=11) and HRV-A (n=5) (Table S1b). The same microbial species was identified in the NP swab and serum samples in 6 patients, one each with HRV-A, HRV-C, hepatitis A virus, hepatitis B virus, rotavirus A, and parvovirus B19.

Bacteria identified in NP samples included four dominant genera, which together comprised 79% of all microbial reads—*Moraxella* (39.4%), *Haemophilus* (16.7%), *Streptococcus* (16.2%), and *Corynebacterium* (6.6%). Given that diversity loss in the microbial flora in lower respiratory tract samples correlates with pneumonia [14], [15], we compared the Simpsons diversity Index (SDI) in patients with and without clinical diagnoses of respiratory tract infection. We found no significant difference between patients with (mean SDI = 0.51, IQR 0.37 −0.65) or without (mean SDI = 0.51, IQR = 0.42 - 0.65; p = 0.86) diagnoses of respiratory infection (Figure S1). This finding is consistent with a growing body of work demonstrating that microbial composition of the nasopharynx may not correlate well with that of the lower respiratory tract in patients with pneumonia [16]–[18].

### mNGS of stool

Among the 10 stool samples collected and sequenced, pathogens were detected in 9/10 samples. The three most common pathogens identified were rotavirus A (50%), *Cryptosporidium* (40%), and human parechovirus (40%). Seven children had co-infections: two rotavirus A and human parechovirus, and one each rotavirus A and *Cryptosporidium*, rotavirus A and enterovirus, *Cryptosporidium* and human parechovirus, *Cryptosporidium* and Norwalk virus, and *Giardia* and human parechovirus. Five samples also had *Blastocystis hominis*, a microbe of uncertain pathogenicity.

### Orthobunyaviruses

Serum from one patient admitted with a clinical diagnosis of malaria and pneumonia contained a novel orthobunyavirus in addition to *P. falciparum*. Assembly of a draft genome and comparison with existing orthobunyavirus genomes indicated that this draft sequence includes 97.5%, 100% and 91% of the L, M, and S coding regions, respectively (Fig 3). Average read coverage across the segments was 86-fold. Phylogenetic comparison showed that the novel virus was significantly divergent from known orthobunyaviruses, sharing 44.9-55.1% amino acid identity with the closest known relatives, Calchaqui virus, Kaeng Khoi virus, and Anopheles A virus (Figure 3, Figures S3a-c). Of note, despite the divergence of this virus, 47.9% of the reads that belong to this new genome were detectable using default parameters in the IDseq pipeline, allowing for facile subsequent assembly. The virus was isolated from a patient from Nyangole village, Tororo District—hence, we propose the name “Nyangole virus”, consistent with nomenclature guidelines for the family *Bunyaviridae*.

**Figure 3:**
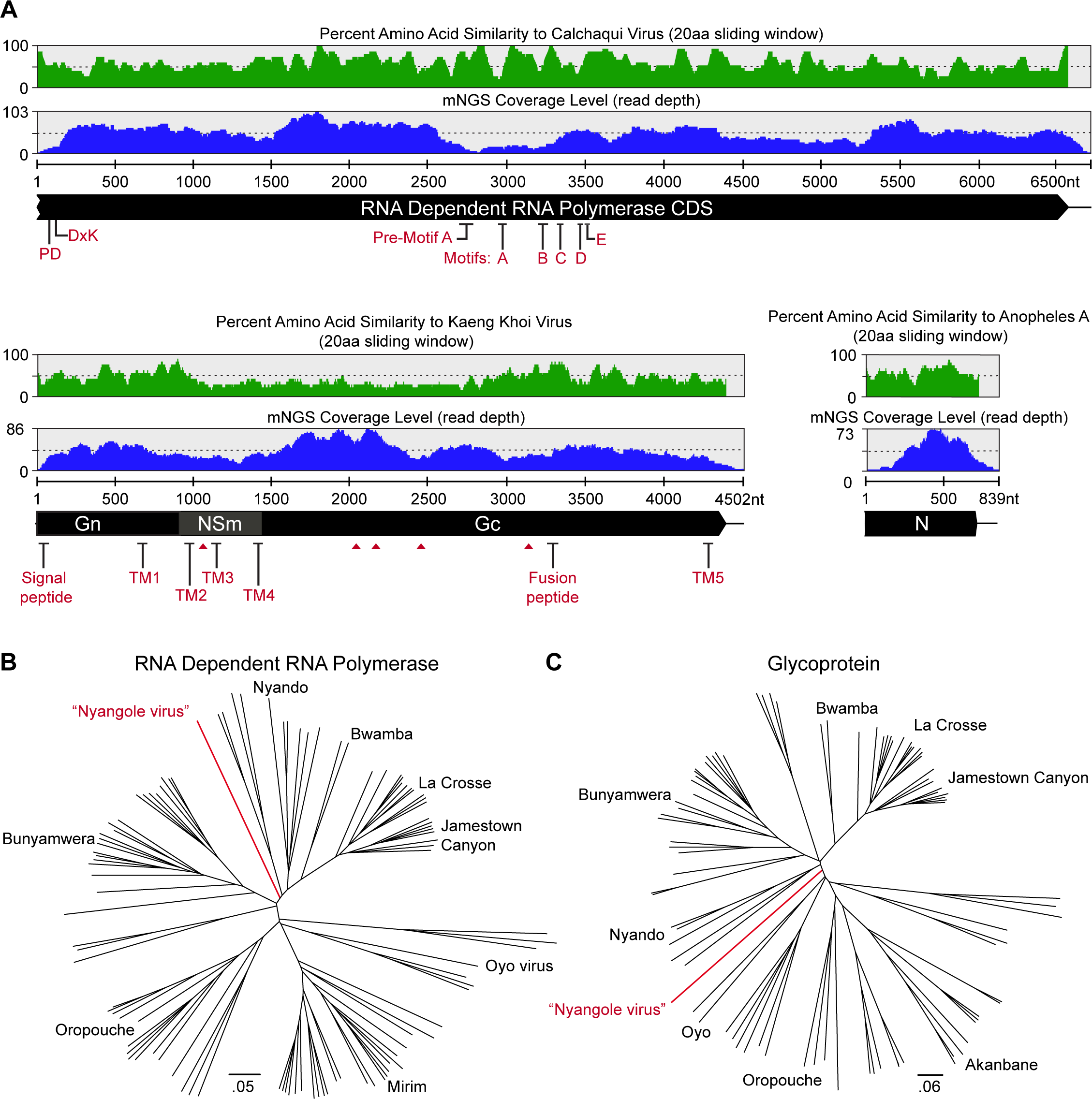
Characterization of the novel orthobunyavirus identified in a febrile child. (A) Schematic representation of the large (L) or RNA dependent RNA polymerase, medium (M) or polyprotein of Gn, NSm and Gc proteins and small (S) segment encoding the nucleocapsid (N) protein of Nyangole virus and percentage identity with the most closely related virus. Phylogenetic tree of all complete orthobunyavirus genome sequences along with Nyangole virus are represented in (B) for the RNA dependent RNA polymerase and (C) for the glycoprotein.

In addition, a second orthobunyavirus, Bwamba virus, was identified in the NP swab sample from a patient admitted with rash, sepsis, and diarrhea. Insufficient sample and sequencing reads precluded genome assembly of this virus.

### Genomic characterization of human rhinoviruses and influenza B in NP swabs

#### Human rhinoviruses

Within the species rhinovirus, we assembled *de novo* a total of 13 HRV-C (mean coverage: 39-fold) and 13 HRV-A (mean coverage: 268-fold) genomes (> 500 bp (Figure 4)). Of these, 10 HRV-A and 9 HRV-C genomes had complete coverage of the VP1 region, which is used to define enterovirus types [19]. HRV types are defined by divergence of >73% in the VP1 gene. As such, we found three HRV-A and eight HRV-C types in this cohort. One individual harbored two distinct HRV-A types (genome pairwise identity=75.3%, VP1 pairwise identity=67.1%). Additionally, we assembled two novel HRV-C isolates from two patients admitted with gastroenteritis (patient ID: EOFI-014), and pneumonia, malaria and diarrhea (patient ID: EOFI-133), that shared 70.1% and 70.7% nucleotide sequence identity at VP1 compared to the closest known HRV-C (Accession JQ245968 and KF688606, respectively). The Picornavirus Working Group has established that novel HRV-Cs should exhibit at least 13% nucleotide sequence divergence in the VP1 gene [20], qualifying these two isolates as novel.

**Figure 4:**
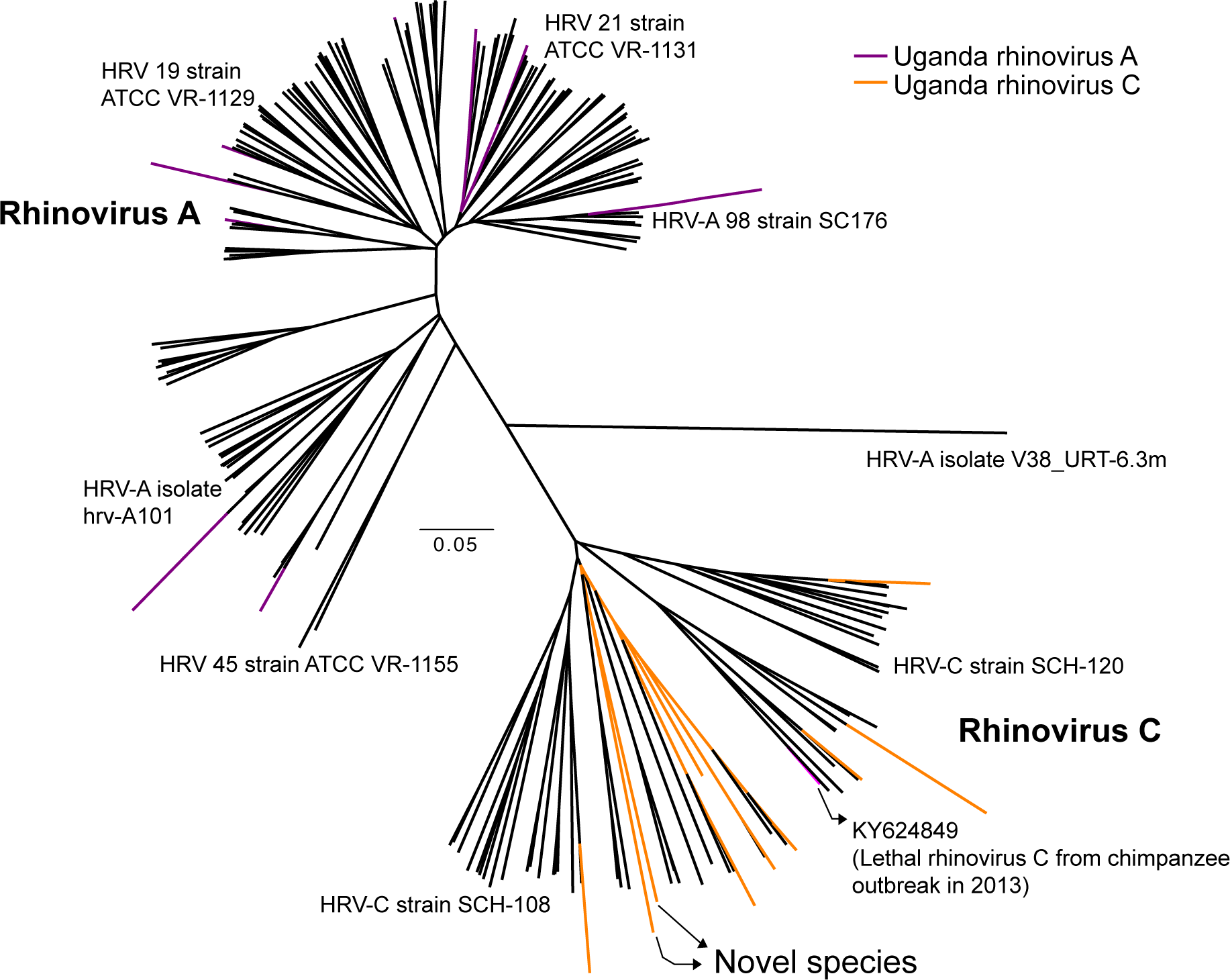
Phylogenetic tree of all complete HRV genomes from NCBI and HRV genomes assembled in this study (Purple – Rhinovirus A, Orange – Rhinovirus C)

#### Influenza B virus

We assembled influenza B genome segments (>500bp, mean-coverage: 41-fold) from six of seven samples containing influenza B virus (one sample had insufficient sequencing reads). The viruses assembled were >99% similar to each other and >99% identical to the B/Massachusetts/02/2012-like virus included in the vaccine recommended by WHO for the 2013-2014 northern hemisphere and 2014 southern hemisphere influenza seasons (Accession numbers: NC_002204 to NC_002211).

#### Antimicrobial resistance profiling in the nasopharynx and stool

We employed Short Read Sequencing Typing (SRST2) to survey the resistance gene landscape of our metagenomic dataset [21]. In total, 86% of patients were found to harbor molecular evidence of resistant organisms in their nasopharynx and 50% in their stool. For both stool and NP swabs, genes conferring resistance to beta lactams were the most abundant, and included *ampC* (n=2), which confers resistance to first through third generation cephalosporins, and *IMP-1* (n=1) which confers broad spectrum resistance that includes carbapenems. In addition, genetic signatures of resistance to other antibacterial agents including aminoglycosides, macrolides/lincosamides, phenicols, sulfas, and tetracyclines were identified (Figure S2).

## Discussion

The clinical management of children with fever is challenging in Africa, where clinicians often have access only to malaria diagnostics. A better understanding of the microbial agents causing fever in African children is needed to inform the development of better diagnostic algorithms and therapeutic guidelines. To address this unmet need, we developed and deployed IDseq in combination with unbiased mNGS to characterize the etiology of fever in Ugandan children admitted to a rural district hospital. We identified a wide range of pathogens in these children.

Other studies evaluating causes of febrile illness in African children have focused on a limited number of pathogens [22]–[25]. In a study of febrile children in Tanzania utilizing rapid diagnostic tests, serologic tests, culture, and molecular tests, viruses accounted for 51% of lower respiratory infections, 78% of systemic infections, and 100% of nasopharyngeal infections. Additionally, 9% of the children had malaria and 4.2% bacteremia [26]. In febrile children in Kenya, reported pathogens were spotted fever group *Rickettsia* (22.4%), influenza (22.4%), adenovirus (10.5%,) parainfluenza virus 1-3 (10.1%), Q fever (8.9%), RSV (5.3%), malaria (5.2%), scrub typhus (3.6%), human metapneumovirus (3.2%), group A *Streptococcus* (2.3)% and typhus group Rickettsiae (1.0%) [27], [28]. Another study reported bacteremia in 19.1% of children admitted to a referral hospital in Uganda [29]. Additionally, in patients (across all age groups) with severe febrile illness, bacteremia was detected in 10.1% in North Africa, 10.4% in East Africa, and 12.4% in West Africa [30].

Traditional pathogen detection methods, including culture, serology, and pathogen directed molecular methods, are logistically challenging in resource limited settings due to the need for extensive microbiology laboratory infrastructure. Unbiased sequencing approaches are designed to identify all potential pathogens, but have been limited by high cost. The costs of deep sequencing are rapidly decreasing, but the analysis of mNGS data necessarily incurs an increasingly large cost, as the available genomic databases to be searched continue to expand. Other major challenges for mNGS approaches include the need for better control datasets, in particular to enable discrimination of contaminants and commensal organisms from true pathogens. The IDseq platform was based on our prior experience with in-house pipelines for pathogen detection [5], [31], [33], and it aims to address existing computational and bioinformatics barriers by providing facile cloud-based mNGS analysis without the requirement for significant on-premise computational infrastructure.

Using mNGS and IDseq, the most common pathogen identified in the blood of febrile Ugandan children was *P. falciparum,* an expected result considering the high incidence of malaria in Tororo District at the time of this study [33]. Some discrepancies were seen compared to blood smear readings, with false positive smears probably due to errors in slide reading, a common problem in African clinics [34], and false negative smears due to the expected greater sensitivity of mNGS for identification of *P. falciparum*. In children with only sub-microscopic parasitemia, it is uncertain whether fevers can be ascribed to malaria, and in fact many children had both *P. falciparum* and additional pathogens identified. Interestingly, three of the four cases of parvovirus B19 were found in association with *P. falciparum*; this co-infection has been associated with severe anemia with life-threatening consequences [35]–[37].

HRV was the most commonly identified virus in NP swab samples, consistent with findings previously reported in sub-Saharan Africa and developed countries [38]–[42]. HRV-C was most frequently encountered (54.1%), followed by HRV-A (43.2%) and HRV-B (2.7%), similar to the distribution of HRVs previously reported in Kenya [38]. We identified two novel HRV-C species; these viruses were about 70% identical to the most closely related previously described HRV-C species [20]. Overall, we detected at least three HRV-A and eight HRV-C types co-circulating in Tororo District. Of note, during the same collection period, a lethal HRV-C outbreak was reported in chimpanzees in Kibale National Park, in western Uganda [43]. The HRV-C reported in western Uganda was modestly related to an isolate observed in our study (74% nucleotide identity; 81% amino acid identity) (Figure 4) [43]. Our results confirm that a wide spectrum of HRVs infects Ugandan children. In addition to HRV, we detected a spectrum of other known respiratory viruses, including RSV, human parainfluenza viruses, human coronaviruses, and adenovirus.

Diarrheal disease is one of the leading causes of death in children in Africa [44]. Approximately 48% of febrile children in our study presented with diarrhea, but due to logistical constraints stool specimens were available for only 10 cases. Rotavirus A, the leading cause of pediatric diarrhea worldwide [45], was the most commonly identified pathogen in this cohort. Rotavirus vaccination, known to be highly effective, is yet to be implemented in Uganda, but the need is clear [45]. In addition to rotavirus A, we detected *Cryptosporidium*, norovirus, *Giardia*, *B. hominis* and several enteroviruses in stool specimens. Enteroviruses, HRV-C, and mamastrovirus were also identified in the serum of three children with clinical diagnoses of gastroenteritis or diarrhea.

Unbiased inspection of microbial sequences from sera revealed a novel member of the orthobunyavirus genus, tentatively named Nyangole virus, which was identified as a co-infection with *P. falciparum* in a child with clinical diagnoses of malaria and pneumonia. The virus was surprisingly divergent from known viruses, with an average amino acid similarity of 51.6% to its nearest known relatives including Calchaqui, Anopheles A and Kaeng Khoi viruses. Mosquitoes have been proposed as a vector for Calchaqui and Anopheles A viruses; Kaeng Khoi virus has been isolated from bedbugs [46]–[48]. Antibodies to these viruses have been detected in human sera, but their role as human pathogens is uncertain [46]–[49]. While the depth of coverage of Nyangole virus sequence in our patient suggests significant viremia, and other orthobunyaviruses are responsible for severe human illnesses (e.g., California encephalitis virus, La Crosse virus, Jamestown Canyon virus, and Cache Valley virus) [50], it is unknown whether the identified virus was responsible for the presenting symptoms.

NP swab analysis identified another Orthobunyavirus, Bwamba virus, in a child admitted with rash, sepsis and diarrhea. This virus has previously been described to cause fever in Uganda [51]. Our identification of two orthobunyaviruses, including one novel virus, in a small sample of febrile Ugandan children suggests that the landscape of previously unidentified viruses that infect African children and potentially cause febrile illness, is significantly under explored.

Molecular evidence of resistance to every available antibiotic except vancomycin in the WHO Essential Medicines List was identified in this study [52]. Three children had genotypic evidence of ESBL producing organisms predicted to be resistant to ceftriaxone, a frontline antibiotic for severe infections in this region [53]. In addition, one of these three children carried IMP-1, which also confers resistance to carbapenems, a class of antibiotics reserved for the most resistant infections.

Our exploratory pilot study had important limitations. First, our samples were not collected randomly, but rather were a convenience sample due to the logistical constraints of our small clinical study; as such the results should not be seen as broadly representative of pathogens infecting Ugandan children. In particular, the lack of identification of bacteremia in study subjects may have been due to a relative paucity of severe illness, compared to that in other studies, in our cohort of admitted children. Second, clinical evaluation of children followed the standards of a rural African hospital, so diagnostic evaluation was limited to physical examination and malaria blood smears. Much more may be learned by linking rigorous clinical evaluation with mNGS results, and thereby more comprehensively assessing associations between clinical syndromes and specific pathogens.

Despite these limitations, our study demonstrates the utility of the mNGS/IDseq platform, and provides an important cross sectional snapshot of causes of fever in African children. Combining unbiased mNGS and the IDseq platform enabled the identification of likely known and novel causes of pediatric illness, and makes available a powerful new open access tool for the characterization of infectious diseases in resource limited settings

### Potential implications

This study provides a snapshot of pediatric fever in Tororo District, Uganda. mNGS permits universal pathogen detection using a single assay, and thus avoids the need for multiple independent, and often costly tests to determine disease etiology. Despite the promise and decreasing costs of this technology, the extensive computational and bioinformatic infrastructure needed to perform analysis of sequencing data remains a major barrier to implementation of mNGS in the developing world. Here we address the need for bioinformatic democratization and computational capacity building with IDseq, an open access platform to bring infectious disease surveillance to regions where it is needed most. As progress is made toward elimination of malaria in sub-Saharan Africa, it will be increasingly important to understand the landscape of pathogens that account for the remaining burden of morbidity and mortality. The use of mNGS can contribute importantly to this understanding, offering unbiased identification of infecting pathogens.

## Materials and Methods

### Enrollment of study subjects

We studied children admitted to Tororo District Hospital, Tororo, Uganda, with febrile illnesses. Potential subjects were identified by clinic staff, who notified study personnel, who subsequently evaluated the children for study eligibility. Inclusion criteria were: 1) age 2-60 months; 2) admission to Tororo District Hospital for acute illness; 3) documentation of axillary temperature >38.0°C on admission or within 24 hours of admission; and 4) provision of informed consent from the parent or guardian for study procedures.

### Study specimens

NP swabs and blood were collected from each enrolled subject within 24 hours of hospital admission. Approximately 5 ml of blood was collected by phlebotomy, the sample was centrifuged at room temperature, and serum was then stored at −80°C. NP swab samples collected with FLOQSwabs™ swabs (COPAN) were placed into cryovials with Trizol (Invitrogen), and stored at −80°C within ~5 min of collection. For subjects with acute diarrhea (≥ 3 loose or watery stools in 24 hours), stool was collected into clean plastic containers and stored at −80°C within ~5 min of collection. Samples were stored at −80°C until shipment on dry ice to UCSF for sequencing.

### Clinical data

Clinical information was obtained from interviews with parents or guardians, with specific data entered onto a standardized case record form that included admission diagnosis and physical examination as well as malaria blood smear results. For malaria diagnosis, thick blood smears were Giemsa stained and evaluated by Tororo District Hospital laboratory personnel following routine standard-of-care practices. No efforts were made to improve on routine practice, so malaria smear readings represent routine standard-of-care rather than optimal quality controlled reads.

### Metagenomic next-generation sequencing (mNGS)

After shipment to University of California, San Francisco, RNA was extracted from clinical samples as well as positive (HeLa cells) and negative (water) controls, and unbiased cDNA libraries were generated using previously described method [54]. Barcoded samples were pooled, size selected (Blue Pippin), and run on an Illumina HiSeq2500 to obtain 135 base pair (bp) paired-end reads

### Bioinformatic analysis and pathogen identification

Microbial pathogens were identified from raw sequencing reads using the IDseq Portal (https://idseq.net), a novel cloud-based, open-source bioinformatics platform designed for detection of microbes from metagenomic data (Figure 1). IDseq scripts and user instructions are available at https://github.com/chanzuckerberg/idseq-dag and the graphical user interface web application for sample upload is available at https://github.com/chanzuckerberg/idseq-web. IDseq is conceptually based on previously implemented platforms [5], [31], [32], but is optimized for scalable Amazon Web Services (AWS) cloud deployment. Bioinformatics data processing jobs are carried out on demand as Docker containers using AWS Batch. Alignments to the National Center for Biotechnology Information (NCBI) database are executed on dedicated auto scaling groups (ASG) of Amazon Elastic Compute Cloud (EC2) instances, with the number of server instances varied with job load. Fast downloads of the NCBI database from the Amazon Simple Storage Service to each new server instance are enabled by the open-source tool s3mi (https://github.com/chanzuckerberg/s3mi). Initial alignment and removal of reads derived from the human genome is performed using the Spliced Transcripts Alignment to a Reference (STAR) algorithm [55]. Low-quality reads, duplicates, and low-complexity reads are then removed using the Paired-Read Iterative Contig Extension (PRICE) computational package [56], the CD-HIT-DUP tool [57], and a filter based on the Lempel-Ziv-Welch (LZW) compression score, respectively. A second round of human read filtering is carried out using bowtie2 [58]. Remaining reads are queried against the most recent version of the NCBI nucleotide and non-redundant protein databases (updated monthly) using GSNAPL and RAPSearch2 [59], [60], respectively. Reads matching GenBank records in the superphylum Deuterostomia are removed, given the high likelihood that such residual reads are of human origin. The relative abundance of microbial taxa is calculated based on reads per million (rpM) mapped at the genus level. To distinguish potential pathogens from ubiquitous environmental agents including commensal flora, a Z-score is calculated for the value observed for each genus relative to a background of healthy and no-template control samples [31]. An overview of this pipeline is represented in Figure 1.

IDseq can process 150 samples at a given time. As of this writing, the current version of the IDseq pipeline (IDseqv1.8) processes fastq files with approximately 70 million reads, typically containing >99% host sequence, in 34 minutes. Run times may vary depending on demand, percentage of non-host sequence, and autoscaling parameters.

For this study we report species greater than 0 rpM and Z-scores detected in the serum, stool, and NP samples. Consistent with previous studies, low levels of “index bleed through” or “barcode hopping” (assignment of sequencing reads to the wrong barcode/index) was observed within the non-templated control samples [61]. To prevent mis-assignment, when a microbe found in more than one sample, it was reported only when present at levels at least four times the level of mis-assigned reads observed in the control samples. Given the extremely high levels of rotavirus found in stool samples, these samples were run in duplicate, and only microbes identified in both samples and present at levels at least four times the number of reads mis-assigned in the control samples were reported. If the reads identified for a given pathogen were not species-specific, we reported the corresponding genus. For NP and stool samples, because the nasopharynx and intestines are normally colonized with commensal bacteria [62]–[65], only non-bacterial species were reported.

### Genome assembly, annotation and phylogenetic analysis

To more comprehensively characterize the genomes of identified microbes PRICE [56] and St. Petersburg genome assembler (SPAdes) [66] were used to *de novo* assemble short read sequences into larger contiguous sequences (contigs). Assembled contigs were queried against the NCBI nt database using BLAST to identify the closest related microbes. GenBank annotation files from genome sequence records corresponding to the highest scoring alignments were used to identify potential features within the *de novo* assembled genomes. Geneious v10.3.2 was used to annotate newly assembled genomes. Reference genomes for multiple sequence alignments and phylogenetic analyses were downloaded from NCBI. Multiple sequence alignments were generated using ClustalW in MEGA v6.0 and the Geneious aligner in Geneious v10.3.2. Neighbor-joining phylogenetic trees were generated using Geneious v10.3.2 and further assessed using FigTree v1.4.3. Annotation of protein domains in the novel orthobunyaviruses was performed using the InterPro webserver [67] as well as direct alignment against previously known orthobunyaviruses. The TOPCONS webserver [68] was used for the identification of transmembrane regions and signal peptides, and the NetNglyc 1.0 Server (http://www.cbs.dtu.dk/services/NetNGlyc/) for the identification of glycosylation sites.

### Evaluation of NP microbiome diversity

We applied SDI to evaluate alpha diversity of microbes identified in NP samples. For this analysis patients were stratified into two categories based on clinical assignment: respiratory infections (admitting diagnosis of pneumonia, respiratory tract infection, or bronchiolitis; n=52) and all other syndromes (n=39); cases with unknown admitting diagnosis were excluded. SDI was calculated in R using the Veganv2.4.4 package on genus-level reads per million values for all microbes, including bacteria. A Wilcox Rank Sum test was used to evaluate differences in SDI between patients in the two categories.

### Antimicrobial resistance gene identification in nasopharynx and stool

The SRST2 computational package was used to identify antimicrobial resistance genes using the Argannot2 database as previously described [21]. We defined extended spectrum β-lactamase (ESBL) as a β-lactamase conferring resistance to the penicillins, first-, second-, and third-generation cephalosporins including the gene classes proposed by Giske et al [69].

## Availability of data and code

All raw data have been deposited under Bioproject ID: PRJNA483304. Assembled genomes can be accessed in GenBank: Accession numbers: MH685676-MH685701, MH685703-MH685719, MH684286-MH684293, MH684298-MH684334. All the raw data, intermediate data and IDseq reports can also be accessed at https://idseq.net [Project ID: Uganda - all - 2]. All IDseq scripts and user instructions are available at https://github.com/chanzuckerberg/idseq-dag and the graphical user interface web application for sample upload is available at https://github.com/chanzuckerberg/idseq-web. While IDseq is under continuous development and improvement, full version control of processing runs and reference databases are standard features.

## Declarations

### List of abbreviations

AWS: Amazon Web Services
ASG: Auto Scaling Groups
BLAST: Basic Local Alignment Search Tool
EC2: Elastic Compute Cloud
ESBL: Extended spectrum β-lactamase
HHV: human herpesvirus
HIV-1: human immunodeficiency 1 virus
HRV: Human Rhinovirus
IQR: Inter-quartile range
L coding region: Large segment coding region (RNA-dependent RNA polymerase)
LZW: Lempel-Ziv-Welch
M coding region: Medium segment coding region (glycoprotein)
MEGA: Molecular Evolutionary Genetics Analysis
mNGS: metagenomic next-generation sequencing
NCBI: National Center for Biotechnology Information
NP swab: Nasopharyngeal swab
nr database: non-redundant database
nt database: nucleotide database
PRICE: Paired-Read Iterative Contig Extension
rpM: reads per million
RSV: respiratory syncytial virus
S coding region: Small segment coding region (nucleocapsid)
SDI: Simpsons diversity Index
SPAdes: St Petersburg genome assembler
SRST2: Short Read Sequencing Typing
STAR: Spliced Transcripts Alignment to a Reference
TTV: torque teno virus
VP1: Capsid protein VP1
WHO: World Health Organization

## Ethics approval and consent to participate

The study protocol was approved by the Uganda National Council of Science and Technology and the Institutional Review Boards of the School of Medicine, Makerere University-College of Health Sciences and the University of California, San Francisco.

## Authors contributions

SN, JH, MK, OB, TR, AM, PR and JLD contributed to experimental design, data acquisition and sample processing. SN, JH, MK, OB, PR, TR, and AM, contributed to patient recruitment and clinical testing. CB, BD, YJ, JS, RE, and JW developed the IDseq pipeline. AR, KK, CL, SN, PR, MRW, and JLD contributed to data analysis. AR, MRW, PR, and JLD contributed to manuscript writing.

## Acknowledgements

We thank the children and families in Tororo District, Uganda, who participated in this study. We thank Eric Chow, Jessica Lund, and the UCSF Center for Advanced Technology for assistance with sequencing. We recognize Amy Kistler. for intellectual discussions and comments on the paper. We thank Mark Stenglein for his helpful discussions and advice.

## Supplemental figures

**Figure S1:**
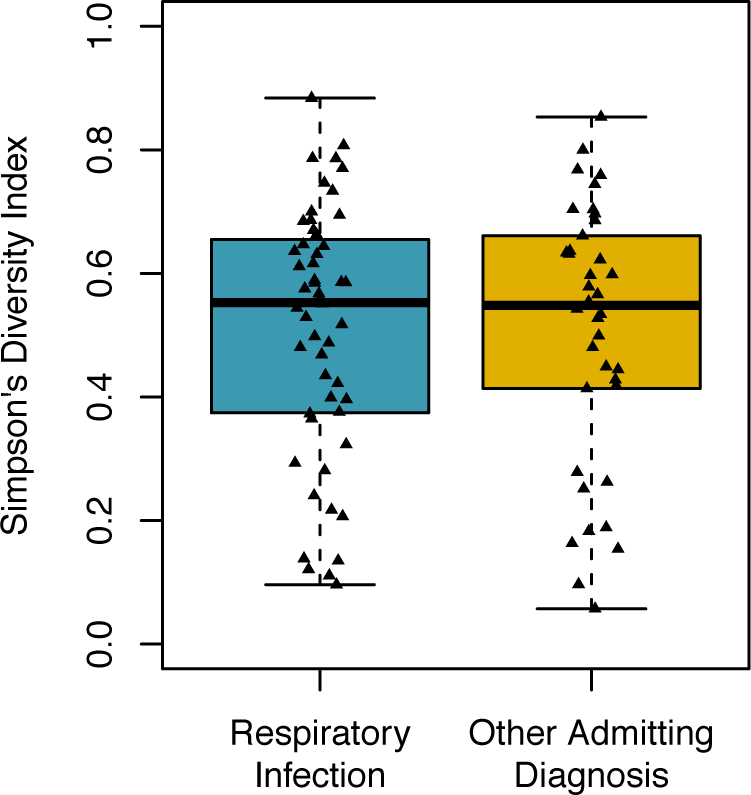
Simpsons diversity index (SDI) for samples with pneumonia versus other etiologies. Each triangle represents one sample.

**Figure S2:**
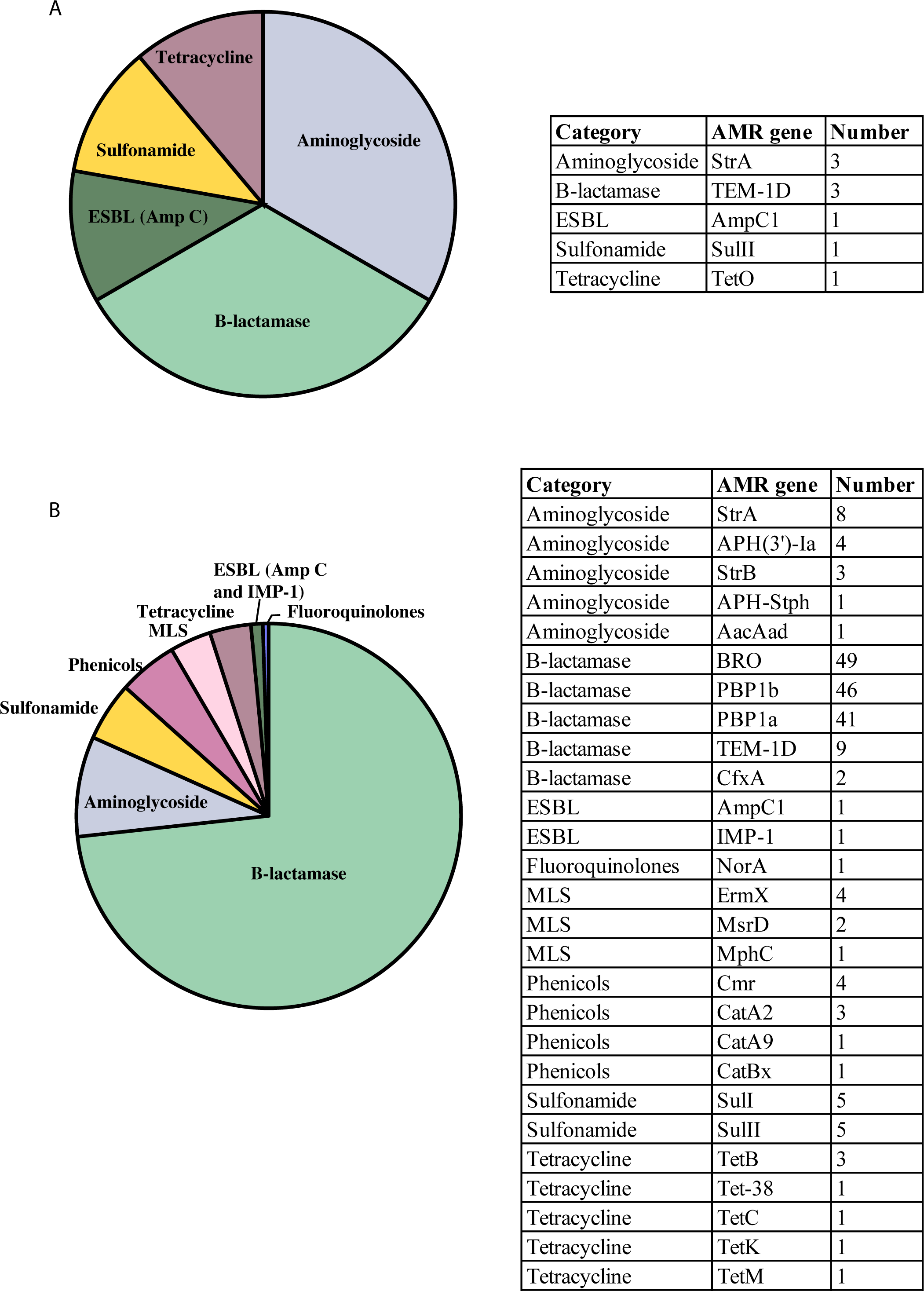
Antimicrobial resistant genes identified in (A) nasopharynx and (B) stool samples

**Figure S3:**
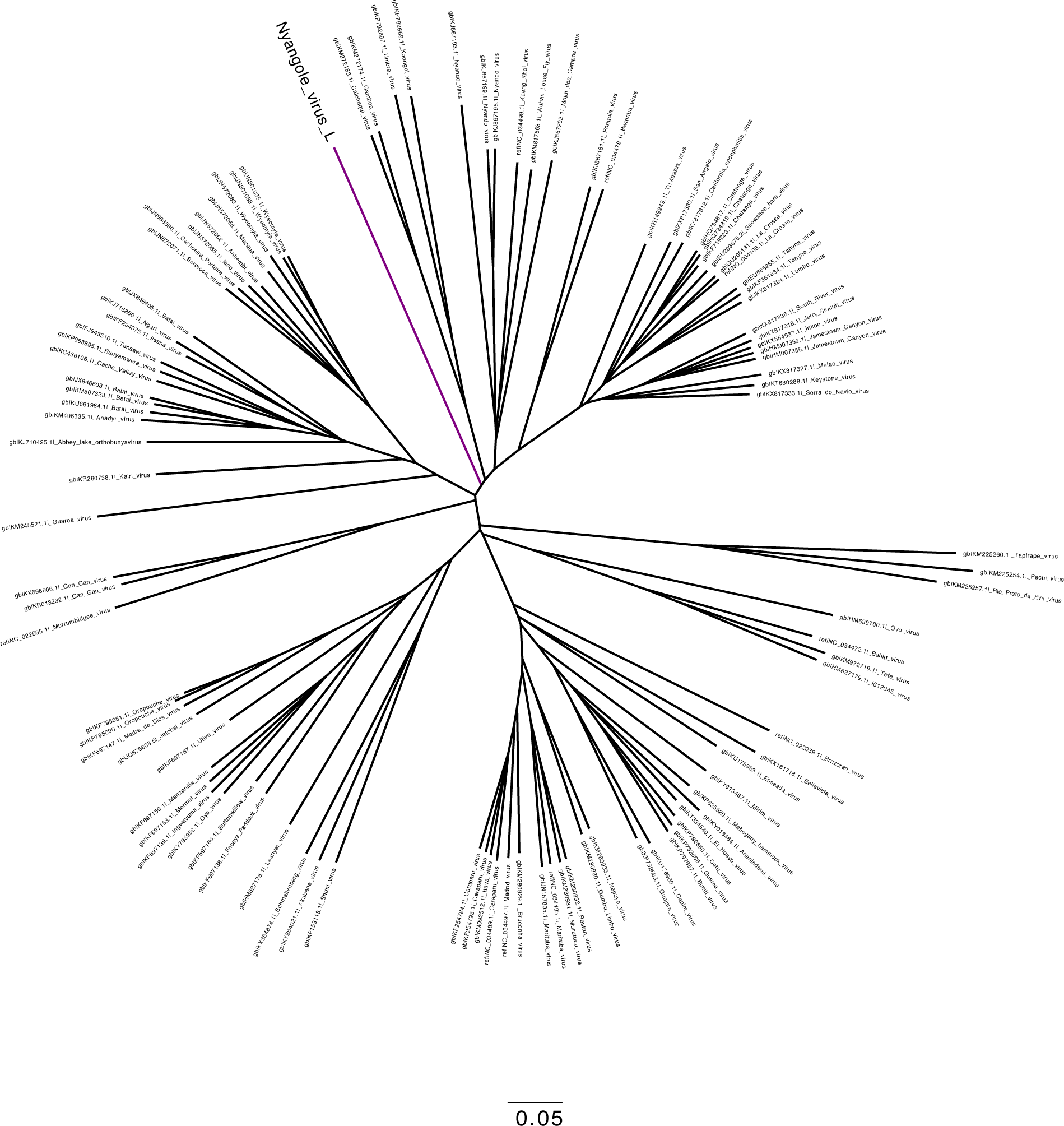
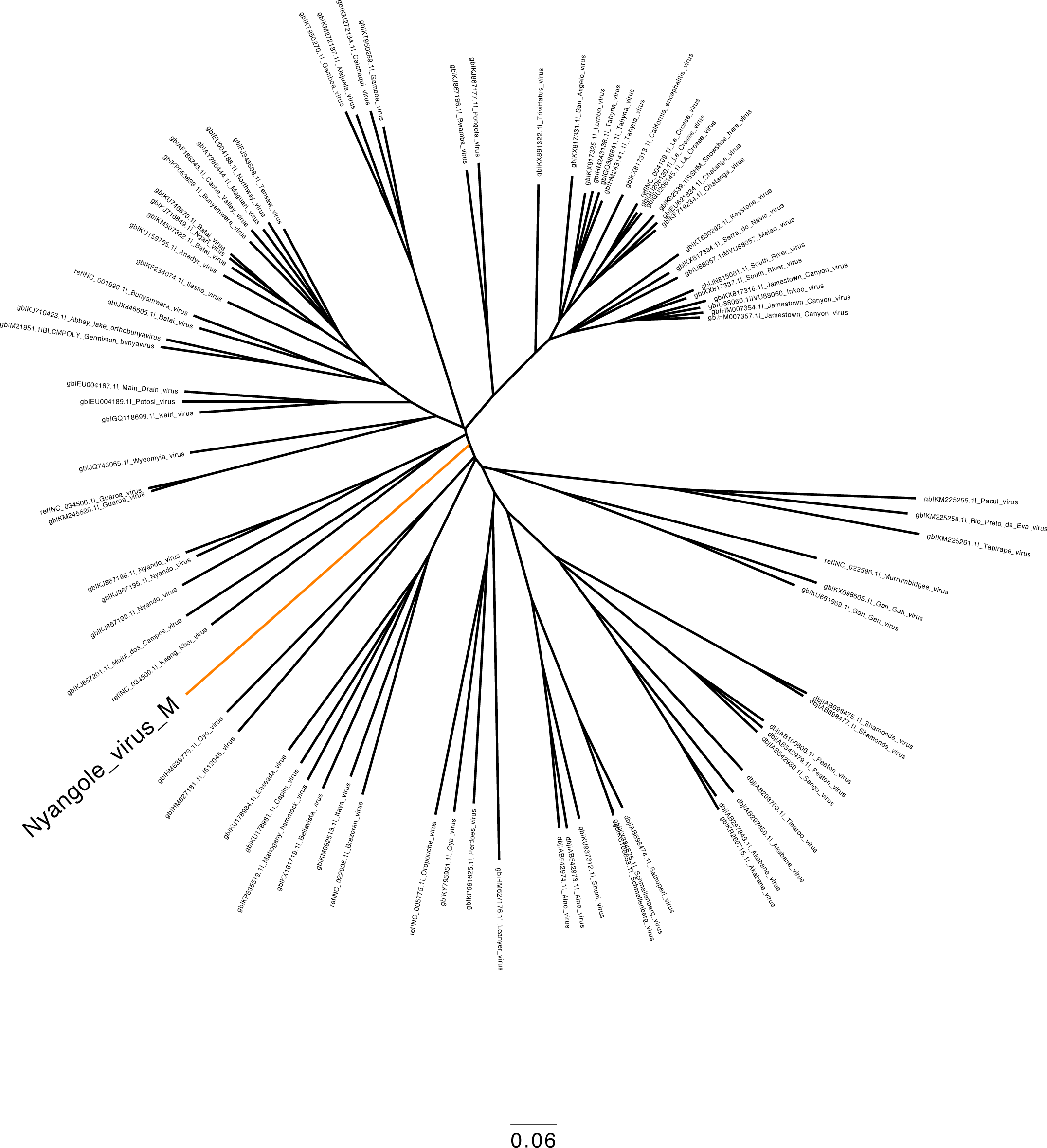
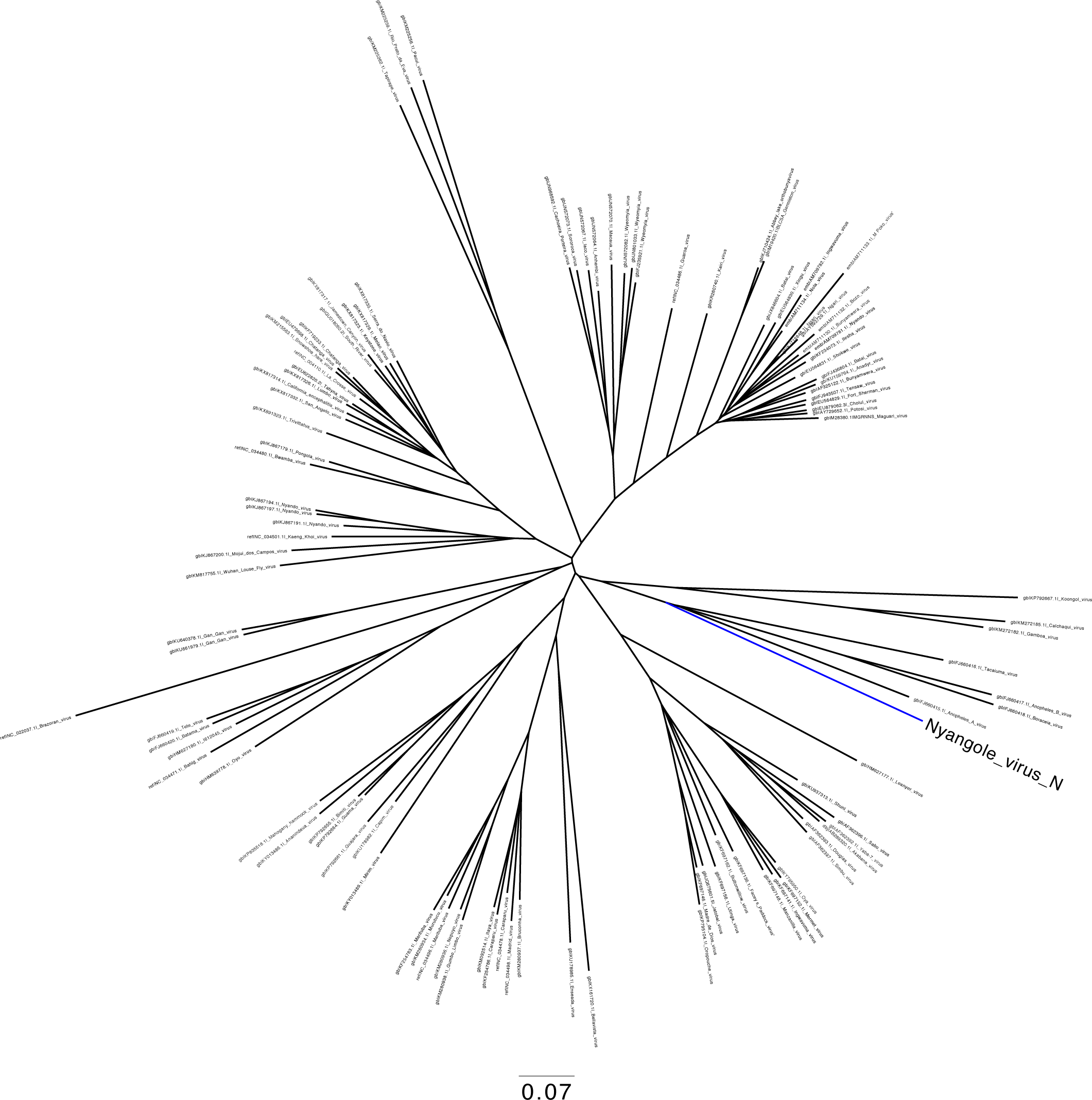
Complete phylogenetic tree of (A) Large (L) or RNA dependent RNA polymerase, (B) Medium (M) or polyprotein of Gn, NSm and Gc proteins and (C) Small (S) segment encoding Nucleocapsid segments

**Table S1:** (A) Co-infection table for *P. falciparum*

| Microbial co-infections with *P.falciparum* | Number of cases |
| --- | --- |
| Parvovirus B19 | 2 |
| Human herpesvirus 4 | 1 |
| Human immunodeficiency virus 1 | 1 |
| Norwalk virus | 1 |
| Orthobunyavirus | 1 |
| HRV-A | 1 |
| HRV-C | 1 |
| Rotavirus A | 1 |
| Parvovirus B19, HRV-C and human parechovirus 2 | 1 |

**Table S1:** B) Co-infection table for HRV

| microbial co-infections with HRV | number of cases |
| --- | --- |
| Human coronavirus OC43 | 2 |
| Human parainfluenza virus 1 | 2 |
| Rotavirus A | 2 |
| Hepatitis A | 1 |
| HHV type 5 | 1 |
| HHV type 6 | 1 |
| Human parainfluenza virus 4 | 1 |
| RSV | 1 |
| KI polyomavirus | 1 |
| RSV and Mamastrovirus 1 | 1 |
| RSV and human parainfluenza virus 1 | 1 |
| HHV type 5 HHV type 7, human coronavirus OC43 | 1 |
| Hepatitis B, human coronavirus NL63 and influenza A | 1 |

## Data files

Data file 1: (A) Clinical signs and symptoms of patients enrolled in the study. (B) mNGS findings in patients enrolled in the study

Data file 2: Total number of sequencing reads and unique non-human reads in all samples analyzed

